# Malignant ascites enhance γδ T cell cytotoxicity towards ovarian cancer via modulating chemokines secretion from the cancer cells that recruits γδ T cells

**DOI:** 10.1101/2025.03.05.641778

**Authors:** Zhanqun Yang, Ying Liu, Mengzhu Zheng, Hui Li, Ruoyao Cui, Pan Wang, Tianhui He, Hongyan Guo, Yinglin Zhou, Jian Lin, Long Chen

**Affiliations:** Department of Pharmacy, Peking University Third Hospital Cancer Center, Peking University Third Hospital, Beijing 100191, China; Beijing National Laboratory for Molecular Sciences (BNLMS), Key Laboratory of Bioorganic Chemistry and Molecular Engineering of Ministry of Education, College of Chemistry and Molecular Engineering, Peking University, Beijing 100871, China; Key Laboratory of Tropical Biological Resources of Ministry of Education, Song Li’s Academician Workstation of Hainan University, School of Pharmaceutical Sciences, Hainan University, Haikou 572000, China; Department of Obstetrics and Gynecology, Peking University Third Hospital, Beijing 100191, China; Synthetic and Functional Biomolecules Center, Peking University, Beijing 100871, China; College of Life Science, Anhui Medical University, Hefei 230032, China

**Author notes:** These authors contributed equally: Zhanqun Yang, Ying Liu, Mengzhu Zheng, Hui Li.

**Keywords:** Ovarian cancer, malignant ascites, γδ T cells, enhance cytotoxicity, chemokines, metabolites

## Abstract

Ovarian cancer patients usually develops peritoneal metastasis and malignant ascites in the advanced stages, which form immuno-suppressive tumor microenvironments that limit the efficacy of immuno-therapies. However, during our previous research trying to develop a γδ T cell-based cell therapy, we noticed that the malignant ascites may enhance the cytotoxicity of γδ T cells towards ovarian cancer cells. Herein, in this work we showed that the phenomenon is real and the low molecular weight components in the ascites act on the cancer to promote the killing by γδ T cells. Transcriptome analysis and in vitro experiments revealed that the malignant ascites induce the secretion of chemokines CXCL2 and CXCL8 by ovarian cancer cells, which recruits γδ T cells through the chemokine receptors CXCR1 and CXCR2, to enhance the cytotoxicity of γδ T cells. Metabolomics analysis discovered compounds that are responsible for the enhancement of γδ T cell cytotoxicity, one of which follows the aforementioned mechanism, while other compounds reflect undiscovered mechanisms. Overall, we presented the positive side of the malignant ascites in anti-tumor immunity, revealed the underlining mechanisms and at least partially interpreted the molecular basis. Our work thus provides new insights into the development of cell therapies for ovarian cancer.

## 1. INTRODUCTION

Ovarian cancer is the six most common and the most lethal gynecological cancer^1^, with a 5-year survival rate of less than 50% and accounts for 5% overall death of cancers in females^2,3^. The standard-of-care treatment for ovarian cancer include chemotherapies, represented by platinum-based drugs^4^, poly (ADP-ribose) polymerase inhibitors (PARPi) and immune checkpoint inhibitors^5–7^. Ovarian cancer is often referred to as the “silent killer” of females due to the lack of early symptoms of the disease that prevent the early diagnosis of ovarian cancer^8^. Therefore, 70% of sovarian cancer patients have progressed into the advanced stages upon diagnosis^9^. One of the main clinical characteristics of advanced stages ovarian cancer patients is the widespread dissemination and implantation of tumor lesions within the abdominal cavity and the continuously formation of large volume of malignant ascites^10^. The malignant ascites not only restrict the efficacy of the first-line chemotherapies and facilitate metastasis^11,12^, but also facilitate the formation of an immune-suppressive tumor microenvironment in the abdominal cavity^13^, which may limit the efficacy of immune-therapies, for example the immune checkpoint blockage therapies^14^. Overall, the formation of malignant ascites is not only an indicator of ovarian cancer progression, but also leads to poor patients prognosis via limiting the efficacy of therapeutics and promoting disease recurrence^12^, suggesting an urgent need for the development of new therapeutics for ovarian cancer with malignant ascites formation.

The adoptive cell transfer therapies, represented by the chimeric antigen receptor T cells (CAR-T) and TCR-Engineered T cells (TCR-T), have achieved tremendous success for the treatment of cancers, especially for the liquid tumors^15–18^. However, the current cell therapies still face great challenges, including limited efficacy to solid tumors^19^, like ovarian cancer, and the necessity of autologous immune cells (i.e., the T cells) due to immunogenicity of the allogeneic immune cells^20^. γδ T cells are a unique subset of T cells^21,22^, which have been reported to be more resistant to the immune-suppressive solid tumor microenvironment and could achieve allogeneic transfer due to the absence of graft-versus-host disease (GvHD) resulted from γδ T cells’ major histocompatibility complex (MHC)-independent antigen recognition property^23^. These properties make γδ T cell a good choice for the development of off-the-shelf allogeneic therapies for solid tumors^24,25^. However, little is known about the effects of the ovarian cancer malignant ascites on the activity of γδ T cells.

Here, in this work, we collected malignant ascites from different ovarian cancer patients and proved that all the ascites samples, and specifically the low molecular weight components act on the cancer cells, enhanced the cytotoxicity γδ T cell towards ovarian cancer cells. Transcriptome analysis and in vitro experiment validations revealed that the malignant ascites induced ovarian cancer cells secretion of chemokines CXCL2 and CXCL8, which in turn recruited γδ T cells to the close proximity of cancer cells that led to the enhancement of γδ T cell cytotoxicity. With the help of untargeted metabolomics, we also discovered several compounds in the ascites that facilitated the enhancement of γδ T cell cytotoxicity towards ovarian cancer cells, of which only one compound follow the chemokine-chemokine receptor mechanism, suggesting a more complicated mechanism of the malignant ascites in enhancing γδ T cell cytotoxicity. Our work thus provided new insights for the development of γδ T immune cell-based therapeutics for ovarian cancer with malignant ascites formation.

## 2. RESULTS

### 2.1 Malignant ascites universally enhance γδ T cell cytotoxicity towards ovarian cancer cell lines and primary ovarian cancer cells

During our previous research using γδ T cell to treat ovarian cancer, we accidently found that the malignant ascites from ovarian cancer patients enhanced the cytotoxicity of γδ T cell towards ovarian cancer cells in vitro. To confirm this phenomenon, we collected ascites from different patients and performed cytotoxicity assay in vitro with ovarian cancer cell lines and primary ovarian cancer cells. Ovarian cancer cell line OVACAR-8 was first tested (**Figure 1A**). Results showed that all tested ascites enhanced the cytotoxicity of γδ T cell towards OVCAR-8 in vitro (**Figure 1A**). This phenomenon was further confirmed with another ovarian cancer cell line SK-OV-3 (**Figure 1B**). We also collected four primary ovarian cancer cells and tested the effects of malignant ascites on these cells. Results showed that the cytotoxicity of γδ T cell towards all these four primary ovarian cancer cells were enhanced (**Figure 1C**). Together, these data proved that the malignant ascites universally enhance γδ T cell cytotoxicity towards ovarian cancer cells.

**Figure 1.**
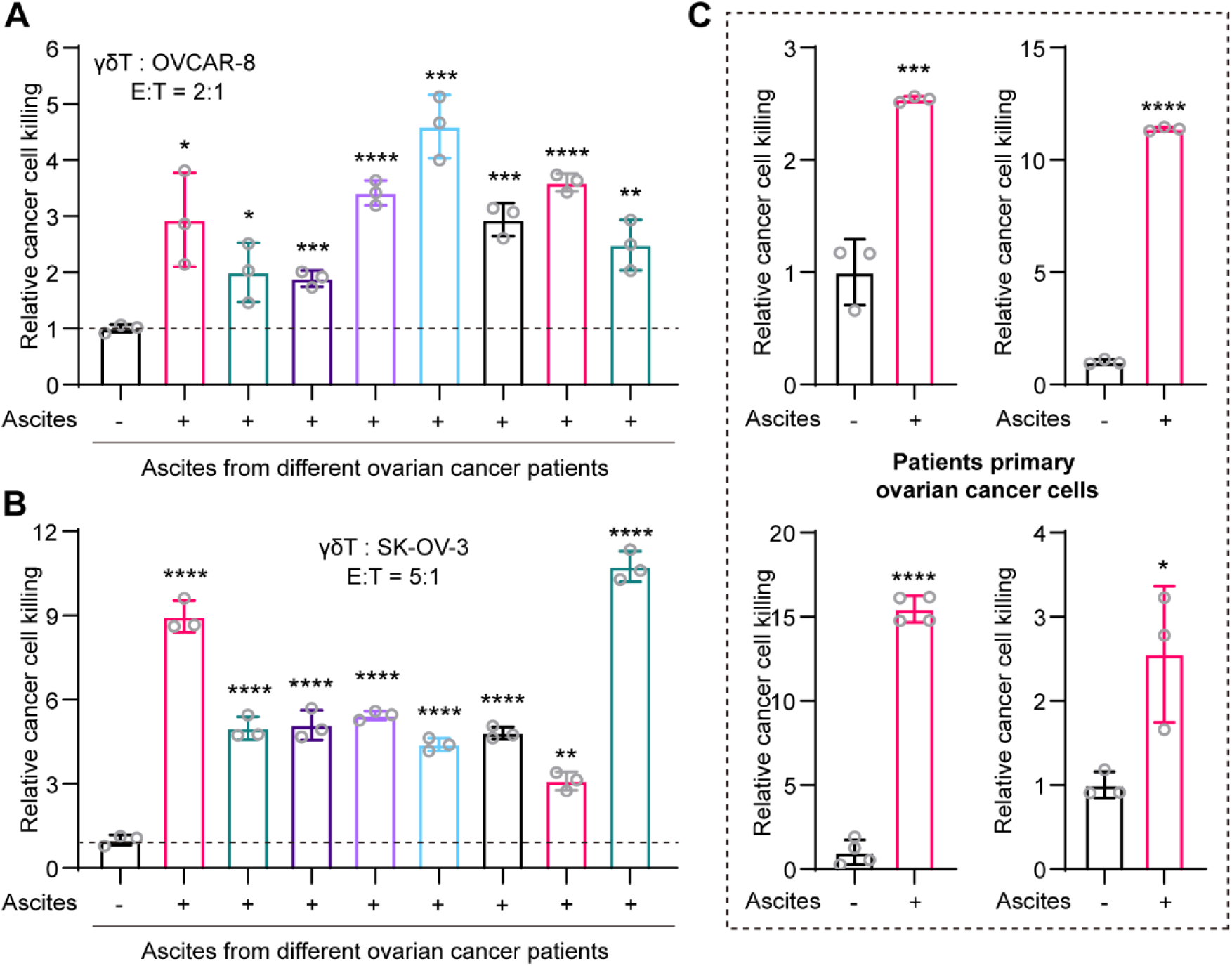
Malignant ascites enhance γδ T cell cytotoxicity towards ovarian cancer cells in vitro. A. Malignant ascites enhance γδ T cell cytotoxicity towards ovarian cancer cell line OVCAR-8 in vitro. B. Malignant ascites enhance γδ T cell cytotoxicity towards ovarian cancer cell line SK-OV-3 in vitro. C. Malignant ascites enhance γδ T cell cytotoxicity towards four patients-derived primary ovarian cancer cells in vitro. For all experiments malignant ascites from different ovarian cancer patients were tested with similar results. Data were presented as mean±SD. n=3 or 4. ns: not significant, * p < 0.05, ** p < 0.01, *** p < 0.001, **** p < 0.0001.

### 2.2 Small molecules from the malignant ascites enhance γδ T cell cytotoxicity towards ovarian cancer cells via acting on cancer cells

As the malignant ascites are complexity mixture, which includes both biomolecules (i.e., proteins) and low molecular weight small molecules^26,27^, we then tried to figure out which component is functional. The low molecular weight small molecules were separated from the mixture using ultrafiltration with a 3 kD cut-off. Cytotoxicity assay showed that the small molecule component also enhanced the cytotoxicity of γδ T cell towards ovarian cancer cell OVCAR-8 (**Figure 2A**). And the results were confirmed with another ovarian cancer cell A2780 (**Figure S1**). While, the remaining high molecular weight components showed greatly reduced or completely abolished enhancement on γδ T cell cytotoxicity (data not shown), indicating that small molecule component of the ascites promote γδ T cell cytotoxicity towards ovarian cancer. We next pretreated the cancer cells or γδ T cells with ascites before the cytotoxicity assay to determine whether the malignant ascites act on the cancer cells or the γδ T cells. Results showed that pretreatment of γδ T cells with different ascites almost had no enhancement on γδ T cell cytotoxicity (**Figure 2B**), while pretreatment of ovarian cancer cells with different ascites led to significant enhancement of γδ T cell cytotoxicity, comparable to co-treatment (**Figure 2B**). Collectively, these results showed that the small molecule components of malignant ascites enhance γδ T cell cytotoxicity towards ovarian cancer cells via acting on cancer cells.

**Figure 2.**
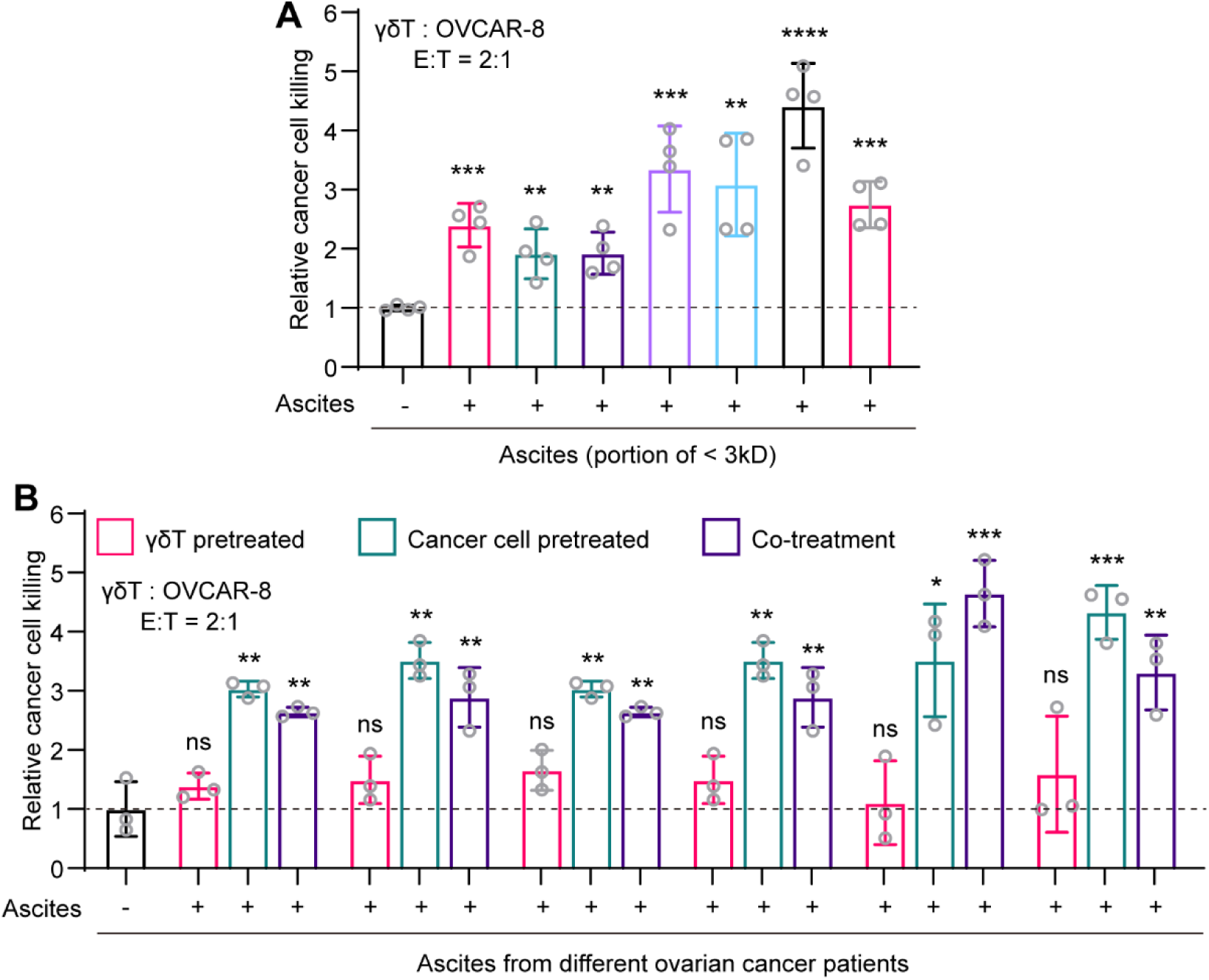
Malignant ascites enhance γδ T cell cytotoxicity towards ovarian cancer cells through the low molecular components via acting on the cancer cells. A. Low molecular weight components of the malignant ascites enhance γδ T cell cytotoxicity towards ovarian cancer cell line in vitro. B. Malignant ascites act on cancer cells to enhance γδ T cell cytotoxicity. For all experiments malignant ascites from different ovarian cancer patients were tested with similar results. Data were presented as mean±SD. n=3 or 4. ns: not significant, * p < 0.05, ** p < 0.01, *** p < 0.001, **** p < 0.0001.

### 2.3 Transcriptome sequencing reveals critical changes in ovarian cancer cells after malignant ascites treatment

To unveil the mechanism underlining the enhancement of γδ T cell cytotoxicity towards ovarian cancer cells by malignant ascites, we performed on transcriptome sequencing using ovarian cancer cell OVCAR-8 before and after treatment with ascites for 3 hours. Volcano plot indicated critical transcriptional changes after ascites treatment (**Figure 3A**), including significant changes of *Cxcl8, Cxcl2, Cxcl1* and *Ccl2.* Kyoto Encyclopedia of Genes and Genomes (KEGG) analysis and Gene Set Enrichment Analysis (GSEA) also revealed signaling pathway enrichment including TNF signaling pathway and cytokine related signaling pathways (**Figure 3B-C**). In consideration of the critical roles of cytokines play in anti-tumor immunity^28,29^, we focused on the cytokines/chemokines CXCL8 (IL-8), CXCL2, CXCL1 and CCL2. Database analysis revealed positive correlation between these cytokines/chemokines expression and patients’ survival in ovarian cancer (**Figure 3D**), indicating beneficial roles of these cytokines/chemokines. Herein, we envisioned that the malignant ascites may induce upregulation of these cytokines/chemokines in ovarian cancer to promote γδ T cell-mediated cytotoxicity.

**Figure 3.**
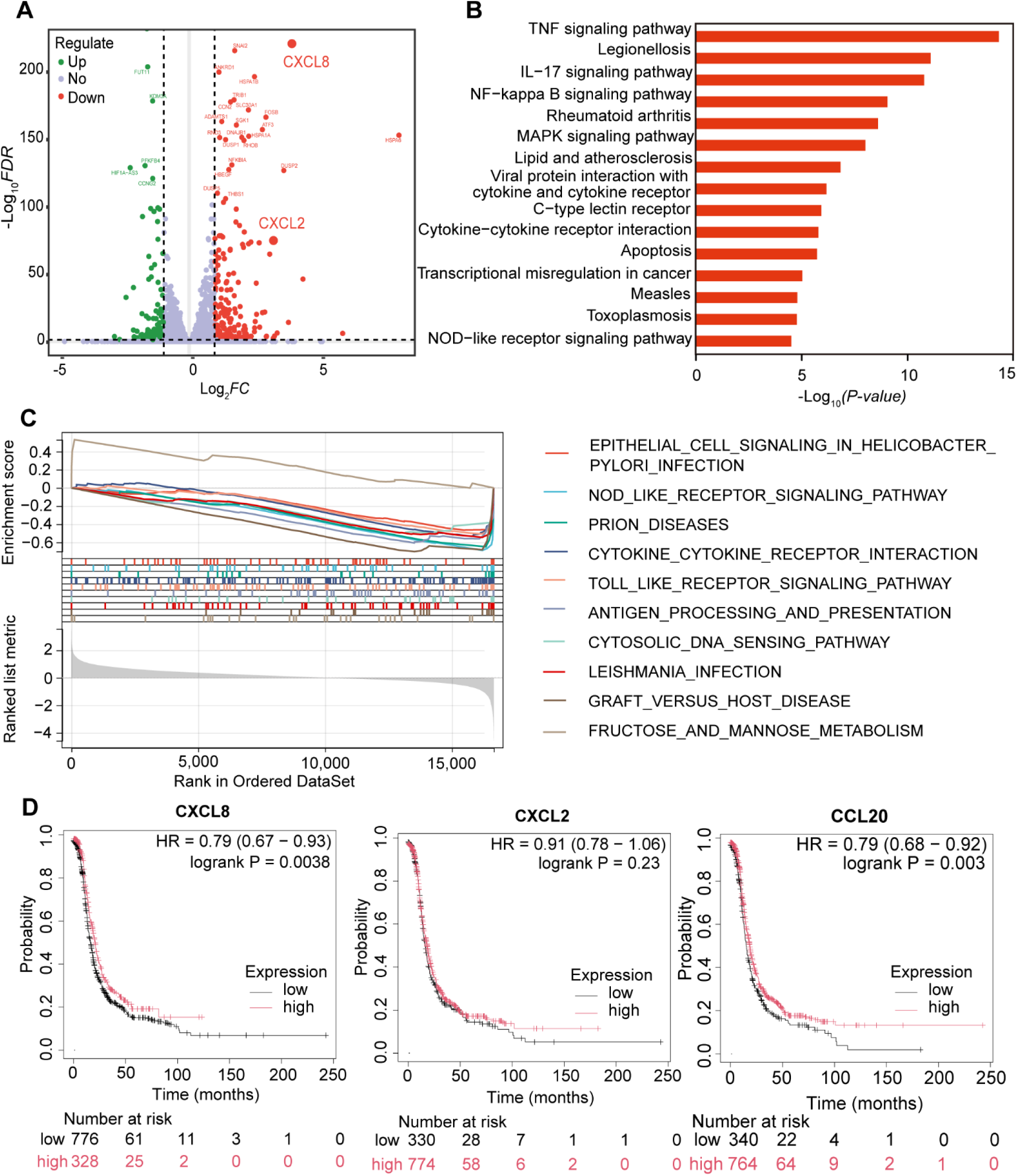
Transcriptome sequencing analysis of ovarian cancer cell treated with malignant ascites. A. Volcano plot showing transcriptional changes of ovarian cancer cell OVCAR-8 after treatment with malignant ascites for 3 hours. B. Kyoto Encyclopedia of Genes and Genomes (KEGG) analysis of the differently expressed genes in OVCAR-8 after ascites treatment. C. Gene Set Enrichment Analysis (GSEA) of the differently expressed genes in OVCAR-8 after ascites treatment. D. Kaplan-Meier survival curves showing the correlation between CXCL8, CXCL2 and CCL20 expression and patients’ survival in ovarian cancer patients.

### 2.4 Validation of the chemokines secretion by ovarian cancer cells after malignant ascites treatment

As the transcriptome sequencing has revealed certain critical changes of chemokines at transcriptional level, we next tried to verify their expression at protein level. Enzyme-linked immunosorbent assay (ELISA) was established to verify the secretion of chemokines CXCL8 (IL-8), CXCL2, CXCL1 and CCL2 in ovarian cancer cell OVCAR-8 after treatment with different malignant ascites. Results showed that 6 out of 7 ascites samples promoted the secretion of CXCL8 by cancer cells (**Figure 4A**), while 5 out of 7 ascites samples promoted the secretion of CXCL2 by cancer cells (**Figure 4B**). For the 7 tested ascites samples, at least one chemokine (CXCL8 or CXCL2) secretion was upregulated by the ascites treatment (**Figure 4A-B**). Meanwhile, validation of either CCL2 or CXCL1 showed no uniformed enhancement of secretion by the ascites (only 3 ascites samples promote CCL20 secretion and 2 ascites samples promote CXCL1 secretion) (**Figure S2A-B**). Collectively, these results revealed that the secretion of CXCL8 and CXCL2 chemokines by ovarian cancer cells were uniformly upregulated by ascites treatment.

**Figure 4.**
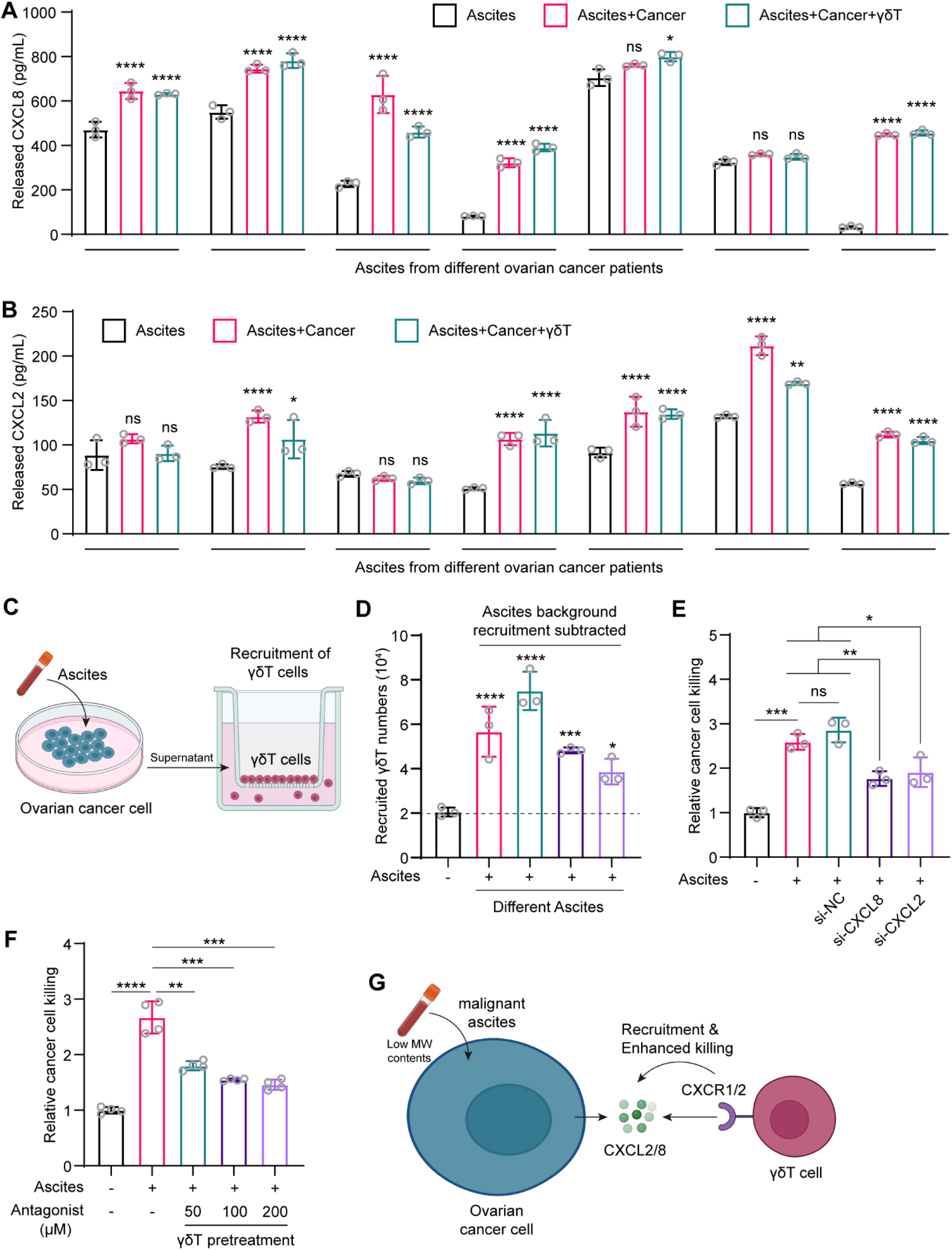
Malignant ascites treatment results in secretion of chemokines CXCL8 and CXCL2 that recruit γδ T cell to enhance its cytotoxicity towards cancer cells. A. ELISA validation of CXCL8 secretion from ovarian cancer cells treated with malignant ascites from different patients for 3 hours. B. ELISA validation of CXCL2 secretion from ovarian cancer cells treated with malignant ascites from different patients for 3 hours. C. Schematic representation of the γδ T cell recruitment assay using ascites treated ovarian cancer cell supernatant. D. Recruitment of γδ T cells by cell culture medium supernatant of ovarian cancer cell treated with ascites from different patients. The background recruitment of the ascites themselves were subtracted. E. Knockdown of CXCL8 and CXCL2 attenuates the effects of ascites in enhancing γδ T cell cytotoxicity towards ovarian cancer cells. F. Pretreatment of γδ T cells with CXCR1/2 antagonist attenuates the effects of ascites in enhancing γδ T cell cytotoxicity towards ovarian cancer cells. G. Proposed mechanism that malignant ascites promote ovarian cancer cell secretion of chemokines CXCL2 and CXCL8 which recruit γδ T cell via the chemokine receptor CXCR1 and CXCR2 to enhance the cytotoxicity of γδ T cell towards cancer cells. Data were presented as mean±SD. n=3 or 4. ns: not significant, * p < 0.05, ** p < 0.01, *** p < 0.001, **** p < 0.0001.

### 2.5 Secretion of chemokines CXCL8 and CXCL2 recruits γδ T cell to cancer cells via the chemokine receptors CXCR1/2 to promote γδ T cell cytotoxicity towards cancer cells

Chemokines are known to recruit certain immune cells to the inflammation sites through the chemokine receptors expressed on immune cells, we thus proposed that the enhanced γδ T cell cytotoxicity towards cancer cells after ascites treatment may result from enhanced recruitment of γδ T cells to cancer cells by the upregulated secretion of chemokines CXCL8 and CXCL2 by cancer cells. To verify this hypothesis, we set up a transwell-based γδ T cells recruitment assay (**Figure 4C**). Culture medium supernatant from ovarian cancer cells with or without ascites treatment was added to the lower well to recruit γδ T cells. To avoid interference of the ascites itself, the background recruitment by ascites was subtracted. The results showed that treatment of the cancer cells with all the 4 tested ascites samples led to enhanced recruitment of γδ T cells (**Figure 4D**), supporting that the enhanced γδ T cell cytotoxicity was a result of close proximity between γδ T and cancer cells. The enhanced recruitment of γδ T cells to the cancer cells were further visualized in the coculture assay under microscope, showing that many more γδ T cells were recruited to the bottom of the culture plate on where the cancer cells residence (**Figure S3**).

We further set up assays to confirm the functions of both the secreted chemokines by cancer cells and chemokine receptors on γδ T cells. Either *Cxcl8* or *Cxcl2* was knocked down in cancer cells using small interfering RNA (siRNAs) (**Figure S4**). Results showed that knockdown of either *Cxcl8* or *Cxcl2* resulted in reduced γδ T cells cytotoxicity enhancement by ascites treatment (**Figure 4E**). CXCL8 and CXCL2 bind to chemokines receptors CXCR1 and CXCR2^30,31^. Database analysis showed that both the two chemokine receptors are highly expressed on γδ T cells (**Figure S5**). Pretreatment of γδ T cells with CXCR1/2 antagonist^32^ showed concentration-dependent of attenuation of ascites induced enhancement of γδ T cells cytotoxicity towards cancer cells (**Figure 4F**), supporting the critical roles of the two chemokine receptors in mediating ascites-based promotion of γδ T cells cytotoxicity. Collectively, we proposed the mechanism that malignant ascites treatment first induces secretion of chemokines CXCL2 and CXCL8 by ovarian cancer cells, which recruits γδ T cells to the cancer cells through interacting with the chemokine receptors CXCR1/2 expressed on γδ T cells that promotes the cytotoxicity of γδ T cells towards ovarian cancer cells (**Figure 4G**). Although the malignant ascites themselves contain CXCL2 and CXCL8, we envisioned that the enhanced secretion of CXCL2 and CXCL8 from the cancer cells may form a concentration gradient of these two chemokines, which gradually recruits the γδ T cells.

### 2.6 Metabolomics-based strategy to identify molecules that enhance the cytotoxicity of γδ T cell towards ovarian cancer cells

With the above mechanism unveiled (**Figure 4G**), we proposed to identify the key components in the malignant ascites that drive its function of enhancing γδ T cell-mediated cytotoxicity towards ovarian cancer cells. As we have proved that the low molecular weight small molecules drive this function (**Figure 2A and Figure S1**), we therefore performed untargeted metabolomics to identify key small molecule metabolites (**Figure 5A**). Malignant ascites from different ovarian cancer patients were first precipitated using methanol before being subjected to LC-MS/MS analysis. After that the detected ions were mapped to the human metabolites databases to identify potential metabolites in the ascites. Two analysis strategies were adopted to discover the key metabolites that drove the enhancement of γδ T cells cytotoxicity toward cancer cells (**Figure 5A**). For the first strategy, the ascites samples were divided into two groups based on their enhancement efficacies, i.e., high or relative low enhancement on γδ T cells cytotoxicity, with a screening set and a validation set. Mass ion peak aera, representing the relative concentration of one compound, was set as the key parameter that affected enhancement on γδ T cells cytotoxicity. Metabolites that were both screened and validated in the screening and validation sets were considered as potential key metabolite compounds. The volcano plots analysis showed that 8 potential compounds were identified (**Figure 5B**). The names and structures are listed in **Figure 5C**. To confirm the functions of these 8 compounds, we performed γδ T cells cytotoxicity assay. As the results showed, treatment with all 8 compounds led to enhanced γδ T cells cytotoxicity towards ovarian cancer (**Figure 5D**), highly supporting our metabolomics strategy. For another analysis strategy of the metabolomics data, direct correlation analysis between the metabolites concentration and γδ T cells cytotoxicity was performed (**Figure 5A**). One compound (**Figure 5E**) showed high correlation efficiency (**Figure 5F**). γδ T cells cytotoxicity assay was performed use this compound and result showed that this identified compound also enhanced γδ T cells cytotoxicity towards ovarian cancer cells (**Figure 5G**). Overall, using untargeted metabolomics approach and different analysis strategies, we identified a series of compounds from the malignant ascites that could enhanced γδ T cells cytotoxicity.

**Figure 5.**
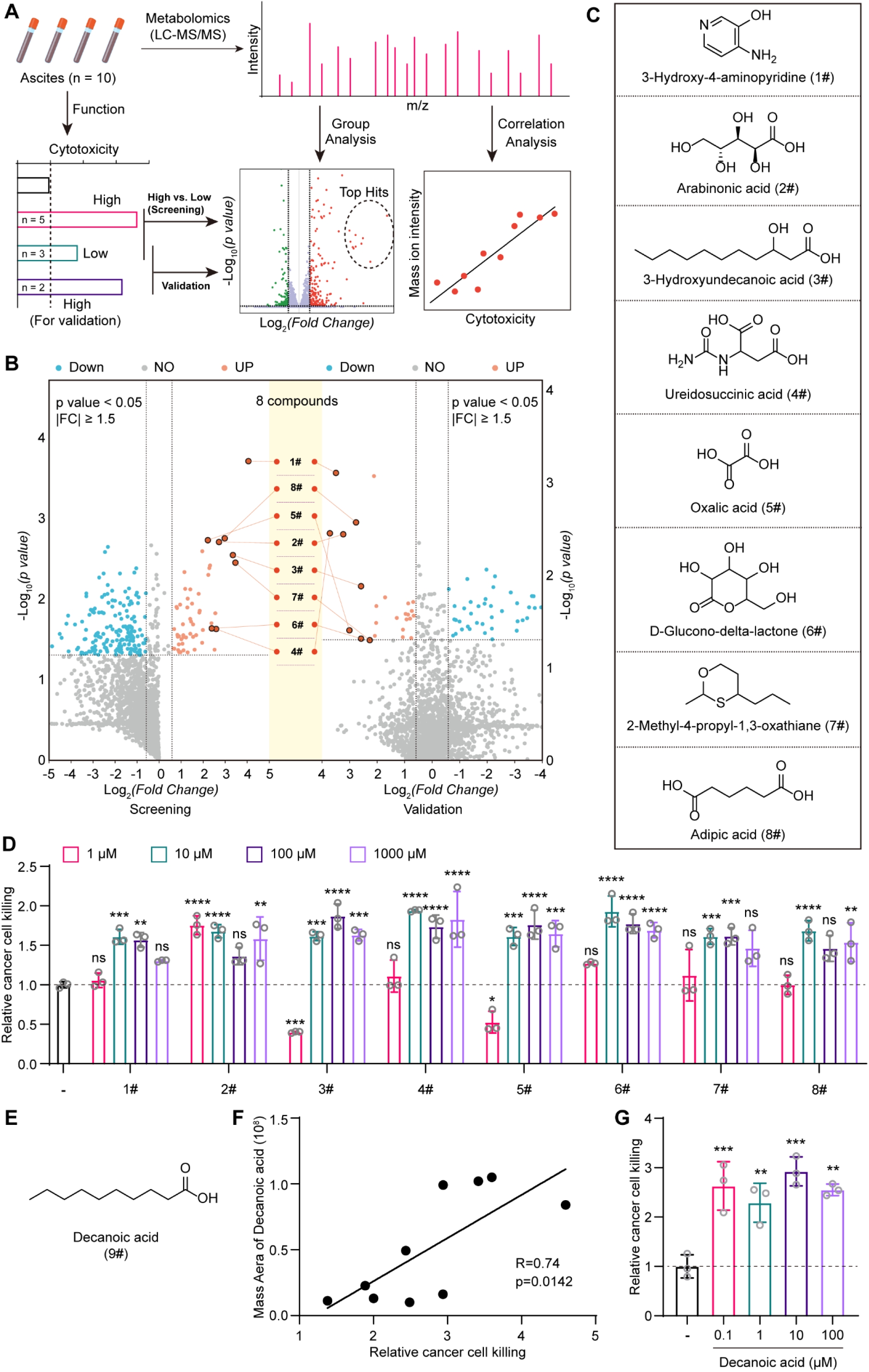
Metabolomics-based strategy to identify molecules that enhance the cytotoxicity of γδ T cell towards cancer cells. A. Schematic showing the metabolomics-based strategy to identify target molecules. For group analysis the ascites from different patients were divided into high and low group based on the enhanced cytotoxicity. For correlation analysis, mass ion intensity of the identified small molecules were correlated to the cytotoxicity. B. Volcano plot showing the screening and validation results for the group analysis which identifies 8 potential molecules. C. Molecular structure and names of the 8 potential molecules. D. In vitro concentration-dependent validation of the effects of the 8 potential molecules on γδ T cell cytotoxicity towards ovarian cancer cells. E. Molecular structure and names of one compound decanoic acid identified from correlation analysis. F. Correlation analysis result of decanoic acid. G. In vitro concentration-dependent validation of the effects of decanoic acid on γδ T cell cytotoxicity towards ovarian cancer cells. Data were presented as mean±SD. n=3. ns: not significant, * p < 0.05, ** p < 0.01, *** p < 0.001, **** p < 0.0001.

### 2.7 Mechanism validation of the identified compounds

We have showed that the malignant ascites drive the enhancement of γδ T cells cytotoxicity through inducing the secretion of CXCL2 and CXCL8 by cancer cells (**Figure 4G**). To further confirm that the identified compounds, we performed ELISA analysis to discover whether these compounds also induce the secretion of CXCL2 and CXCL8 by ovarian cancer cells. Results showed that only compound #5 (oxalic acid) induced enhanced secretion of both CXCL2 (**Figure S6**) and CXCL8 (**Figure 6A**), comparable to the ascites induced enhancement. The malignant ascites are complexity mixture, therefore it’s quite reasonable that the ascites may drive the enhancement of γδ T cells cytotoxicity through multiple mechanisms, one of which is the induction of the chemokines CXCL2 and CXCL8 that recruit the γδ T cells. Herein, we proposed the mechanism that compound #5 (oxalic acid) drives the enhancement of γδ T cells cytotoxicity through the chemokine-chemokine receptor pathway, while other 8 compounds follow other mechanisms that further need to be discovered (**Figure 6B**).

**Figure 6.**
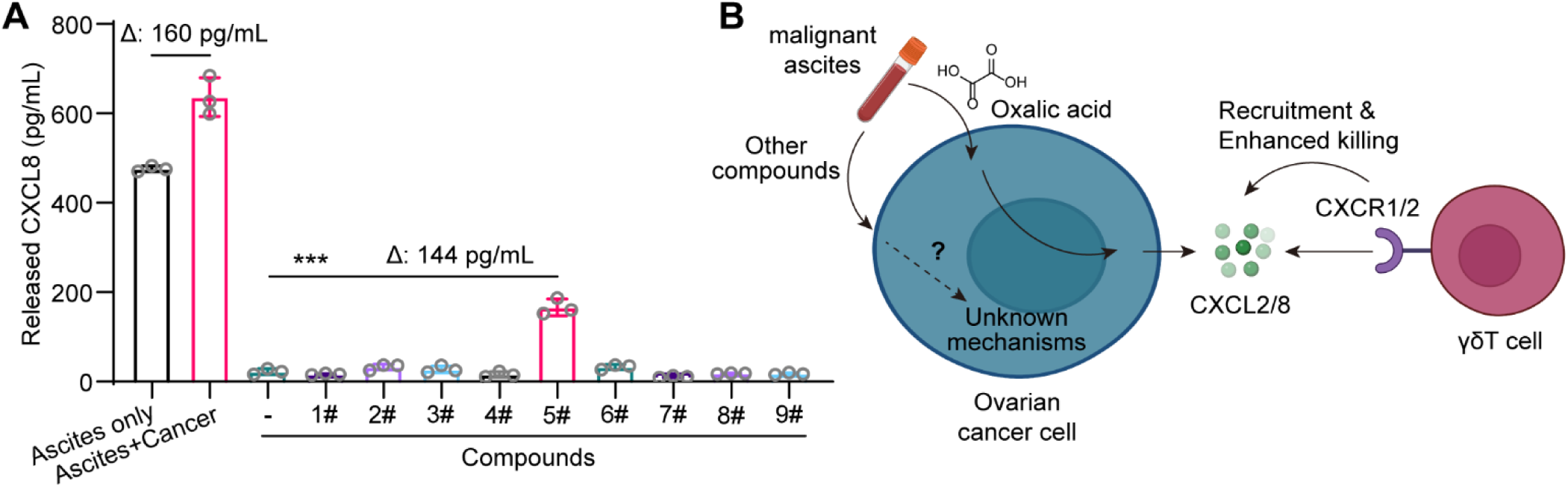
Mechanism validation of the identified molecules. A. ELISA validation of CXCL8 secretion from ovarian cancer cells treated with the 9 identified compounds for 3 hours. Data were presented as mean±SD. n=3. *** p < 0.001. B. Proposed mechanism of the ascites derived compounds on the cytotoxicity of γδ T cell towards cancer cells. Oxalic acid enhance γδ T cell cytotoxicity via promotion of chemokines CXCL2 and CXCL8 secretion from cancer cells that recruit γδ T cells, while the mechanisms for other compounds remain to be unveiled.

### 2.8 Malignant ascites enhance the cytotoxicity of the “CAR-T like” antibody-γδ T cell conjugate towards ovarian cancer

CAR-T (Chimeric Antigen Receptor T-Cell Immunotherapy) has revolutionized the treatment for cancers, especially for hematological cancers^33^. However, besides the genetically engineered CAR-T, chemical engineered T cells have also showed similar clinical potentials^34–39^. In our previous trail, we constructed “CAR-T like” antibody-γδ T cell conjugates (γδ T-ACC) by metabolic glycan labeling and click chemistry (**Figure 7A**), which use the antibody moiety to target cancer cells that leads to the “CAR-T like” enhanced cytotoxicity towards cancer cells (**Figure 7B**) (manuscript under review in another journal and a PD-L1 targeting γδ T-ACC indicated for the treatment of ovarian cancer peritoneal metastasis is under pre-clinical evaluation). We therefore wonder whether the malignant ascites also promote the cytotoxicity of this PD-L1 targeting γδ T-ACC. In vitro experiment revealed that, as expected, malignant ascites from different ovarian cancer patients enhanced the killing of ovarian cancer cells by γδ T-ACC (**Figure 7C**), suggesting that for ovarian cancer patients with malignant ascites formation, the therapeutic outcomes of γδ T-ACC maybe even better.

**Figure 7.**
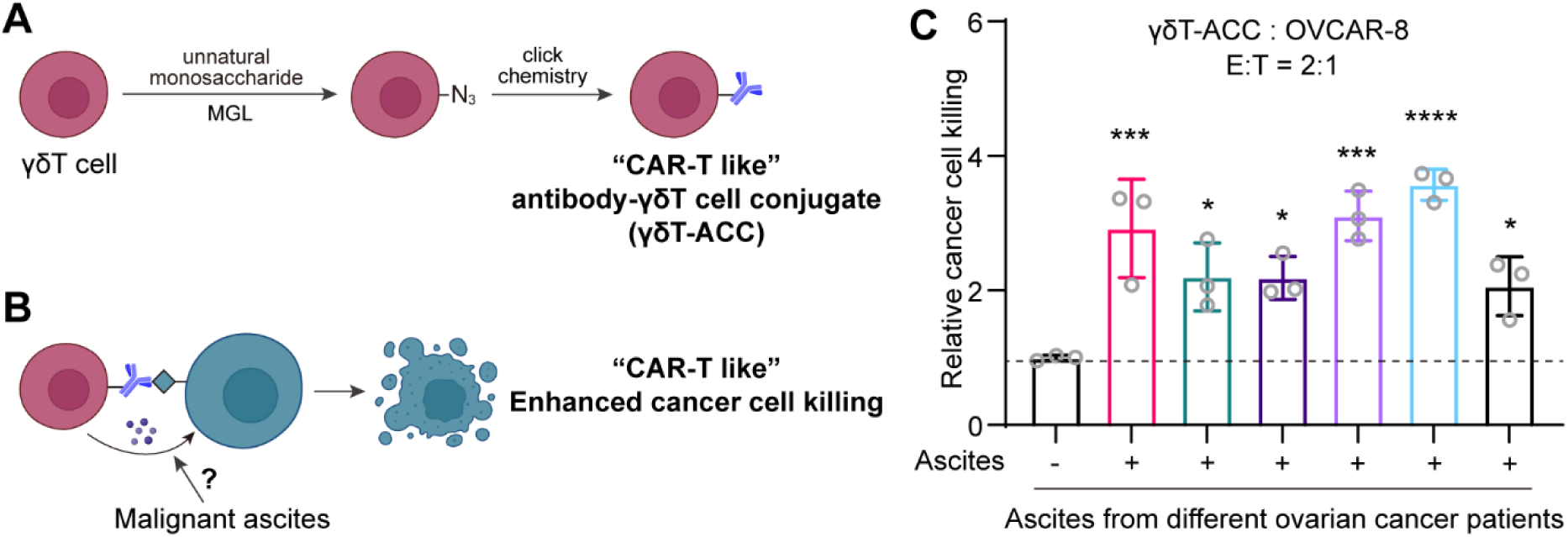
Malignant ascites enhance the cytotoxicity of “CAR-T like” antibody-γδ T cell conjugates. A. Schematic showing the preparation of the “CAR-T like” antibody-γδ T cell conjugates (γδ T-ACC). B. Functional revealing of γδ T-ACC’s enhanced cytotoxicity towards cancer cells with the effects of malignant ascites on the cytotoxicity of γδ T-ACC not being investigated. C. Malignant ascites enhance PD-L1 targeting γδ T-ACC cytotoxicity towards ovarian cancer cell line OVCAR-8 in vitro. Data were presented as mean±SD. n=3. * p < 0.05, *** p < 0.001, **** p < 0.0001.

## 3. DISCUSSION

Ovarian cancer is one of the most common and most lethal cancer in females^1^. Due to the lack of effective early diagnosis methods, over 70% of ovarian cancer patients have developed into the late stages upon their first diagnosis, which leads to the poor prognosis of ovarian cancer^8,9^. Besides, patients in the late stages normally develop peritoneal metastasis lesions with the formation of large volume of malignant ascites^10^. The malignant ascites not only restrict the efficacy of the chemotherapies^11^, but also lead to the formation of a more immune-suppressive tumor microenvironment in the abdominal cavity^13^.

However, in this work we find the second face of the malignant ascites, in which it may enhance the therapeutic efficacy of adoptive T cell transfer therapies. We revealed that the malignant ascites, and specifically the low molecular weight components, treatment of the ovarian cancer cells universally promoted the cytotoxicity of γδ T cells towards ovarian cancer cells (**Figure 1-2**). Transcriptome sequencing reveals that malignant ascites upregulated the expression of *Cxcl2* and *Cxcl8* and ultimately induced the secretion of chemokines CXCL2 and CXCL8 by cancer cells (**Figure 3-4**). The secreted chemokines then recruited the γδ T cells to the close proximity of the cancer cells via interacting with chemokine receptors CXCR1 and CXCR2 highly expressed on γδ T cells (**Figure 4C-G**), which triggered the enhanced cytotoxicity of γδ T cells towards the ovarian cancer cells. To our delight, the chemokine receptors CXCR1 and CXCR2 are highly expressed on γδ T cells, medium expressed on the CD8^+^ T cells and very low expressed on Treg cells (**Figure S5**), suggesting that the malignant ascites may also recruit and enhance the cytotoxicity of CD8^+^ T cells towards ovarian cancer, while not recruit and activate the immune-suppressive Treg cells.

With the help of untargeted metabolomics, we further discovered potential small molecule metabolites in the malignant ascites that promote the killing of ovarian cancer cells by γδ T cells (**Figure 5**). However, among the 9 selected metabolite compounds, only one compound oxalic acid follows the aforementioned chemokine-chemokine receptor mechanism (**Figure 6 and Figure S6**), suggesting more complicated mechanism still need to be discovered. Recently, many researches have reported the discovery of small molecule modulators of T cells functions for the enhancement of adoptive T cell transfer therapies like CAR-T cell therapies. Our discovered small molecule metabolites may find further applications in these strategies.

Our work still face some limitations. As we all know that the ovarian cancer ascites also contains large number of cells^26^, including T cells, dendritic cells and etc^10^. How will these cells or immune cells crosstalk with adoptively transferred γδ T cells and affect γδ T cells’ anti-tumor functions is still unknown and not been studied in this work. But faithfully, we envisoned the adoptive γδ T cell transfer can lead to beneficial effects. The reason is that γδ T cells can function as professional antigen-presentation cells^40^. Therefore, the enhanced cytotoxicity of adoptively transferred γδ T cells towards ovarian cancer cells may lead to elevated antigen-presentation by γδ T cells, which would then potentially activate the T cells in the ascites for better anti-tumor immunity. But overall, more comprehensive research integrating the effects of both the metabolites and floating cells in the ovarian cancer malignant ascites would be insightful in the future.

Overall, our work showed another favorable function of the malignant ascites from ovarian cancer, in which the malignant ascites could enhanced the killing of the cancer cells by γδ T cells. This work shed light on future fundamental researches and translational studies of adoptive γδ T cells transfer for the treatment of ovarian cancer with malignant ascites formation, which account for 90% of the patients with advanced stages of ovarian cancer.

## 4. MATERIALS AND METHODS

### 4.1 Collection and storage of patients’ malignant ascites

Ascites samples were collected from patients undergoing surgery for ovarian cancer at the department of Obstetrics and Gynecology Department of Peking University Third Hospital (Beijing, China) and approval by the Ethics Committee of Peking University Third Hospital (ID: IRBO0006761-M2019291). Ascites samples were collected in the operating room and stored in tubes, which were then transported to the laboratory using an ice box. The collected samples were centrifuged at 4°C for 10 minutes at 350g to remove cellular components, followed by a second centrifugation at 2000g for 10 minutes. The supernatant was stored at - 80°C for subsequent experiments.

### 4.2 Expansion of γδ T cells

γδ T cells are expanded from peripheral blood mononuclear cells (PBMCs) of healthy donors (under the guidelines of the approved protocol: ICB00006761-M2023641). PBMCs are isolated from whole blood by density-gradient centrifugation using Lymphocyte Separation Medium (Solarbio, P8610). For the expansion of γδ T cells, PBMCs are first stimulated with 5 μM Zoledronic acid (MCE, HY-13777) for three days, followed by expansion in CTS OpTmizer T-Cell Expansion SFM medium (Gibco, A1048501) containing 1000 IU IL2 (T&L Biotechnology Co., Ltd., GMP-TL906-0050), after which they are cryopreserved in liquid nitrogen for subsequent experimental use.

### 4.3 Cancer cell culture

Human ovarian cancer cells (OVCAR8, SK-OV-3 and A2780) were acquired from ATCC. Cells were maintained in RPMI 1640 supplemented with 10% fetal bovine serum (Gibco, 26010074), 100 U/mL penicillin/streptomycin (Gibco,15140122) in a humidified atmosphere of 5% CO_2,_ 95% air at 37 °C.

### 4.4 In vitro cytotoxicity assay

Cancer cells and their drug-treated counterparts are first incubated with 5μM Calcein-AM at 37°C under 5% CO_2 c_onditions for 30 minutes. Subsequently, the cells are washed three times with PBS buffer. Labeled cancer cells were added to 96-well plate at a density of 2×10^5 cells/mL with a total volume of 50μL per well. Then, 50μL of ascites fluid and effector γδ T cells are added, maintaining an effector-to-target (E:T) ratio of 2:1, and incubated at 37°C for 4 hours. Wells with the addition of 50μL cell culture medium containing 5% FBS served as controls. γδ T cell cytotoxicity to the cancer cells was determined by the release of Calcein-AM. The background release of Calcein-AM from control wells were defined as 0% cytotoxicity and the maximum release of Calcein-AM from 1% Triton X-100 lysed control wells were defined as 100% cytotoxicity. The cytotoxicity is calculated using the formula: Cytotoxicity % = (Experimental group release - Background release) / (Maximum release - Background release) × 100%. The amount of Calcein-AM release was determined by its fluorescence. To determine the involvement of CXCL2/8 and their receptors CXCR1/2, either the cancer cells were treated siRNA targeting CXCL2/8 (siRNA for *Cxcl2:*′*- GCAUCGCCCAUGGUUAAGAAAdTdT*-3′, 5′-*UUUCUUAACCAUGGGCGAUGCdTdT*-3′; siRNA for *Cxcl8:* 5′-CUUAGAUGUCAGUGCAUAA-3′, 5′-UUAUGCACUGACAUCUAAG-3′; *siNC:* 5′- *UUCUCCGAACGUGUCACGUdTdT*-3′, 5′-*ACGUGACACGUUCGGAGAAdTdT*-3′) or the γδ T cells were pretreated with CXCR1/2 antagonist for 30 minutes at concentration of 50, 100 and 200 μM (Navarixin, catalog # HY-10198, MCE) before subjected to cytotoxicity assay. To determine the effects of identified metabolites, indicated concentrations of purify compounds were added to the coculture systems.

### 4.5 Transcriptome sequencing

OVCAR-8 cells were cultured in the presence or absence of malignant ascites for 3 hours at 37 °C. The cells were then collected and total RNA was extracted. Transcriptome sequencing was performed by Berry Genomics Co., Ltd. following standard protocols.

### 4.6 Untargeted Metabolomics

After the ascites samples were collected, low molecular weight small molecules were separated with 3 kDa cut-off centrifugal filter. 10 µl of filtrate was treated with 4-fold volume of cold methanol to extract metabolites and remove the small peptides. Insoluble materials were centrifuged at 14,000 × g for 15 min, and the resulting supernatants were evaporated. Untargeted metabolomics were performed on a Dionex Ultimate 3000 UHPLC (Thermo, USA) coupled with a Q-Exactive mass spectrometer (Thermo, USA). Samples were resuspended in 50 µl of ultrapure water for LC-MS/MS analysis. Compounds were separated on an XBridge Amide column (100 × 4.6 mm, 3.5 μm; Waters, Milford, MA, USA). Buffer A comprised 20 mM ammonium acetate (pH 9.4) in water, and buffer B was acetonitrile (LC-MS grade). Raw data of metabolomic analysis was performed with Compound Discoverer, metabolite structure and function information were searched in the database HMDB (www.hmdb.ca) and KEGG (https://www.kegg.jp).

### 4.7 RT-PCR

Total RNA from the cells was extracted with RNA extraction kit (Catalog#: R1200, Solarbio, Co., Ltd, China) and cDNAs were generated with M-MuLV reverse transcriptase (Catalog#: R1200, Mei5 Biotechnology, Co., Ltd, China). Primers (F: 5’-CTCAAGAATGGGCAGAAAGC-3’, R: 5’-AAACACATTAGGCGCAATCC-3’ for *Cxcl2*, F: 5’-TCTGCAGCTCTGTGTGAAGGT-3’, R: 5’-TGTGTTGGCGCAGTGTGGT-3’ for *Cxcl8*, F: 5’-GTCTCCTCTGACTTCAACAGCG-3’, R 5’-ACCACCCTGTTGCTGTAGCCAA-3’ for *Gapdh*) and ChamQ SYBR qPCR Master Mix (Catalog#: R1200, Mei5 Biotechnology, Co., Ltd, China) were used for RT-PCR to determine the knockdown efficacy of *Cxcl2* and *Cxcl8*. Each sample was tested in triplicate. Glyceraldehyde-3-phosphate dehydrogenase (GAPDH) was used as a normalization control. The conditions for PCR were: 95℃ for 30s, and 40 cycles of 95℃ for 5s and 60℃ for 30s.

### 4.8 Measurement of chemokine release using ELISA

To determine the chemokine release after malignant ascites treatment, OVCAR-8 cells were treated with ascites or small molecule compounds for 3 hours. Supernatants were collected and the release of the chemokines were determined with ELISA kits following the manufacturer’s instructions. The information for the ELISA kits are as follow: CXCL1 (catalog # SEKH-0065, Solarbio), CXCL2 (catalog # SEKH-0066, Solarbio), CXCL8 (catalog # SEKH-0016, Solarbio), CCL20 (catalog # SEKH-0249, Solarbio).

### 4.9 In vitro γδ T cell recruitment assay

Cell culture medium supernatant of OVCAR-8 cells with or without malignant ascites treatment was added to the lower chamber of transwell (Corning, 3421). γδ T cells were added to the upper insert. Cells were incubated at 37℃ for 6h before the recruited γδ T cells being determined by CCK8 (MedChemExpress, HY-K0301).

### 4.10 Preparation of the PD-L1 targeting γδ T-ACC

γδ T cells were seeded in cell culture dish at a density of 5 × 10^4 cells/mL and incubated with azido containing unnatural monosaccharide (100 μM) overnight to produce γδ T-azide. A human PD-L1 binding nanobody was first labeled with the click chemistry group dibenzocyclooctyne using DBCO-PEG4-NHS (MCE, HY-130809) to produce DBCO-Nb-PD-L1. γδ T-azide were then reacted with DBCO-Nb-PD-L1 at 37 ℃ for 30 min. After that the cells were washed three time with PBS. The resulting cells are the PD-L1 targeting γδ T-ACC.

### 4.11 Public Database Analyses

The kmplot.com database was utilized to investigate the correlation between tumor gene expression and patients survival. The platform amalgamates gene expression profiles from databases including GEO, EGA, and TCGA, enabling the assessment of 54,675 genes’ influence on survival in 21 cancer types. Kaplan-Meier survival curves can be generated, and meta-analyses can be conducted to ascertain molecular markers linked to survival outcomes. In our analysis, we focused on high-grade serous ovarian cancer and identified genes of significant interest, thereafter we extracted the pertinent survival data associated with these genetic markers. Subsequent Kaplan-Meier plotting delineated survival disparities based on gene expression, with log-rank tests employed to assess statistical significance. These findings were contextualized for their prognostic and therapeutic implications.

### 4.12 Statistical analyses

Statistical analysis and graphing of the data were performed using Graphpad prism 9.0.1 (La Jolla, CA, USA). Data are presented as mean ± SD. Statistical significance was determined by one-way ANOVA test for variance after Bonferroni correction. Significance was determined as * *p*-value < 0.05, ** *p*-value < 0.01, \*\*\**p* < 0.001, \*\*\*\**p* < 0.0001.

## AUTHOR CONTRIBUTIONS

L.C., J.L., Y.Z. and H.G. designed the experimental plans. Z.Y., Y.L., M.Z., H.L. and R.C. performed the experiments. P.W. and T.H. collected clinical samples. Z.Y. and L.C. analyzed the data and drafted the manuscript. L.C., J.L., Y.Z., and H.G. revised the manuscript.

## ACKNOWLEDGEMENTS

We acknowledge GraphPad Prism 9 and Biorender.com for the help in processing data and creating figures.

## DATA AVAILABILITY STATEMENT

The data are available from the corresponding author on reasonable request.

## ETHICS STATEMENT

Collection of malignant ascites were approval by the Ethics Committee of Peking University Third Hospital (ID: IRBO0006761-M2019291).

## FUNDING INFROMATION

This work was supported by the National Natural Science Foundation of China (No. 82274034) and Peking University Medicine plus X Pilot Program-Platform Construction Project (2024YXXLHPT004).

