## supporting information for "Malignant ascites enhance γδ T cell cytotoxicity towards ovarian cancer via modulating chemokines secretion from the cancer cells that recruits γδ T cells"

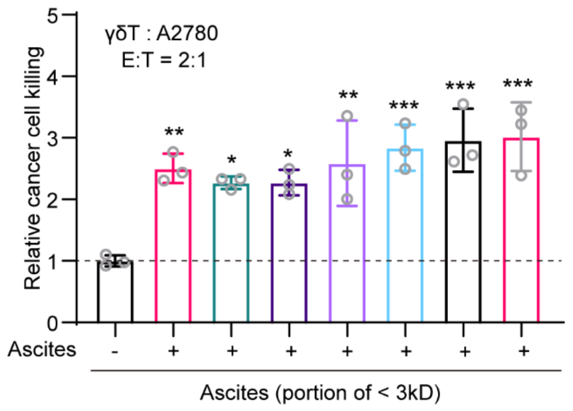


**Figure S1.** Low molecular weight components of the malignant ascites from different patients enhance γδ T cell cytotoxicity towards ovarian cancer cell line A2780 in vitro. Data were presented as mean±SD. n=3. * $p < 0.05$, ** $p < 0.01$, *** $p < 0.001$.

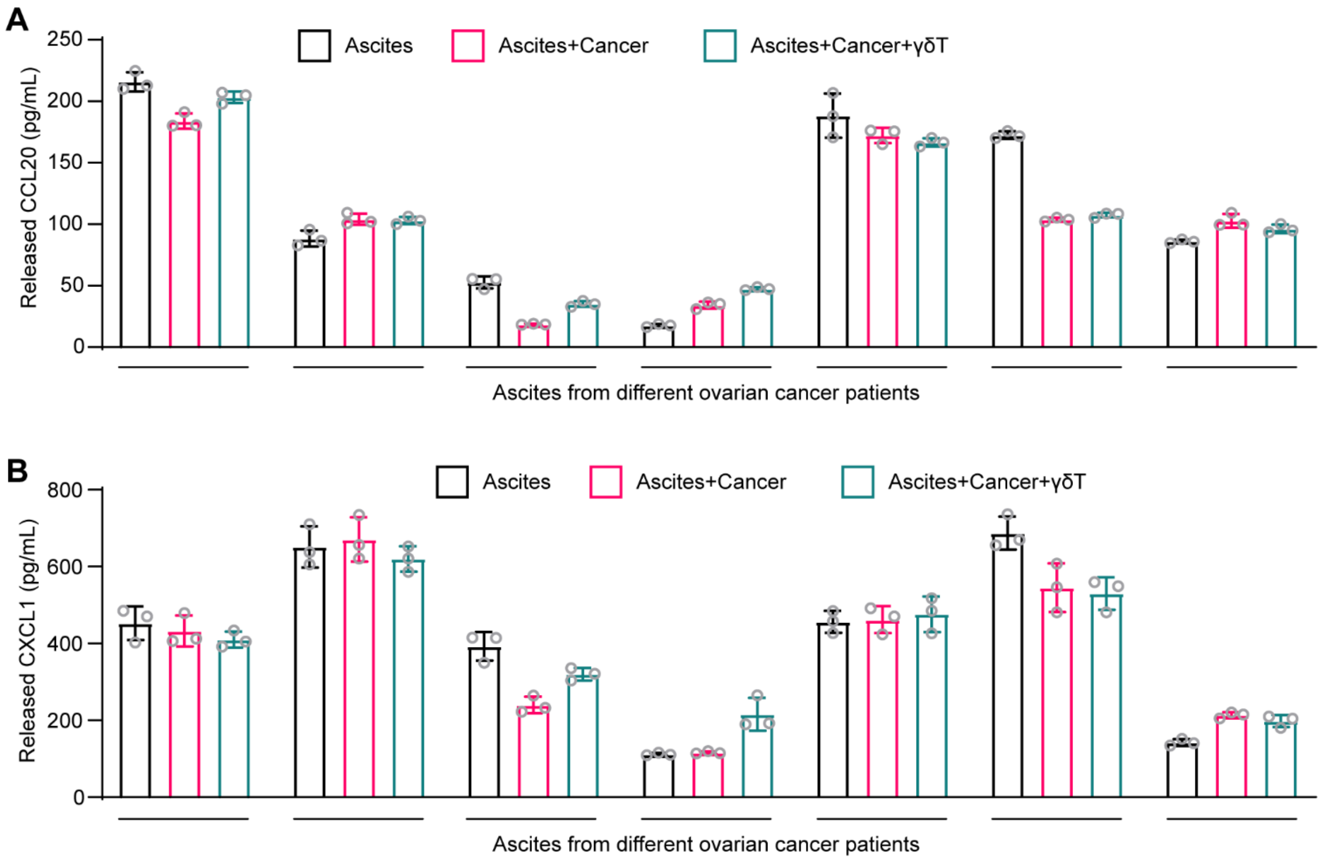


**Figure S2.** A. ELISA validation of CCL20 secretion from ovarian cancer cells treated with malignant ascites from different patients for 3 hours. B. ELISA validation of CXCL1 secretion from ovarian cancer cells treated with malignant ascites from different patients for 3 hours.

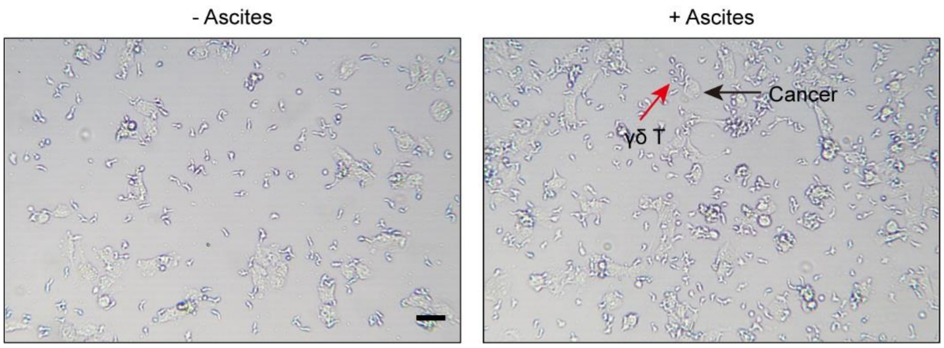


**Figure S3.** Microscope images showing the recruitment of the γδ T cells to the close proximity of the cancer cells on bottom of the cell culture plate. Red arrow indicates γδ T cells and black arrow indicates cancer cells. Scale bar: 50 μm.

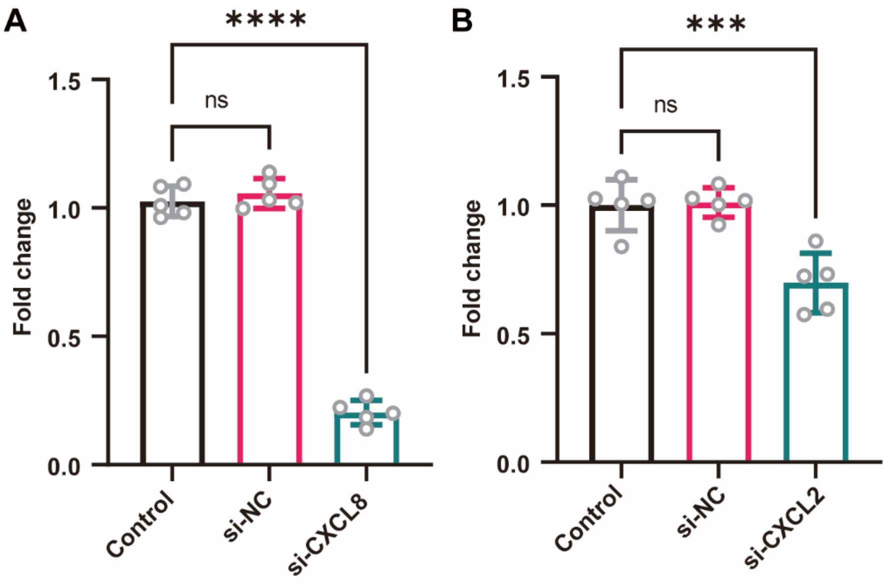


**Figure S4.** RT-PCR analysis of the knockdown of *Cxcl8* (A) and *Cxcl2* (B). Data were presented as mean±SD. n=5. ns: not significant, **** $p < 0.0001$.

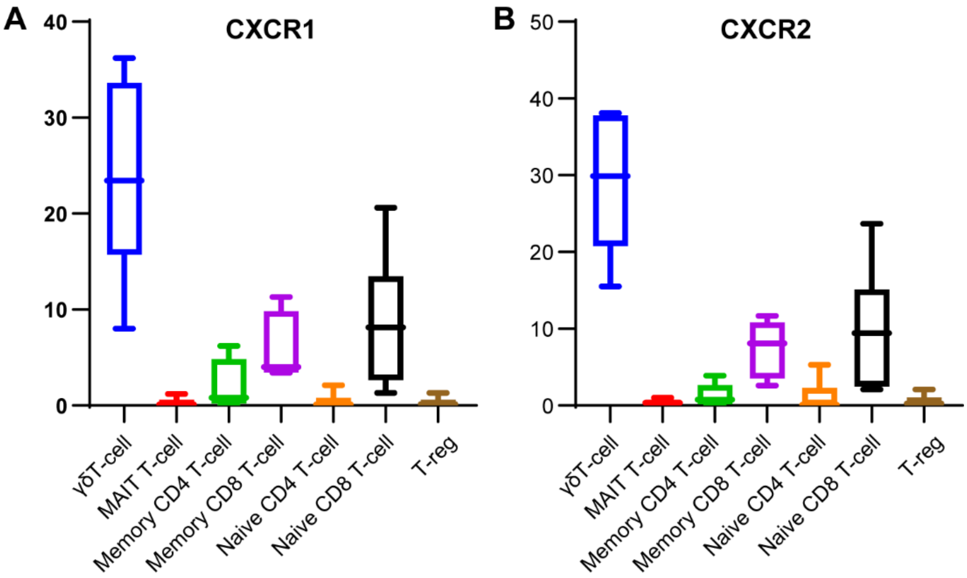


**Figure S5.** CXCR1 (A) and CXCR2 (B) expression in different T cell subtypes. Data acquired from the Human Protein Atlas.

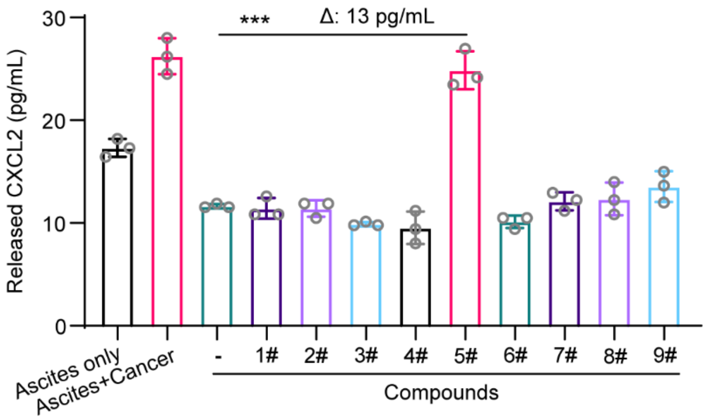


**Figure S6.** MELISA validation of CXCL2 secretion from ovarian cancer cells treated with the 9 identified compounds for 3 hours. Data were presented as mean±SD. n=3. *** $p < 0.001$.
